# Particle interactions and their effect on magnetic particle imaging and spectroscopy

**DOI:** 10.1101/2021.10.29.466424

**Authors:** Lorena Moor, Subas Scheibler, Lukas Gerken, Konrad Scheffler, Florian Thieben, Tobias Knopp, Inge K. Herrmann, Fabian H. L. Starsich

**Affiliations:** Nanoparticle Systems Engineering Laboratory, Department of Mechanical and Process Engineering, ETH Zurich, Sonneggstrasse 3, 8092 Zurich, Switzerland; Particles-Biology Interactions, Department Materials Meet Life, Swiss Federal Laboratories for Materials Science and Technology (Empa), Lerchenfeldstrasse 5, 9014 St. Gallen, Switzerland; Section for Biomedical Imaging, University Medical Center Hamburg-Eppendorf, Lottestraße 55, 22529 Hamburg, Germany; Institute for Biomedical Imaging, Hamburg University of Technology, Am Schwarzenberg-Campus 3, 21073 Hamburg, Germany

**Author notes:** Corresponding author: Fabian H. L. Starsich.

**Keywords:** Magnetic particle imaging, magnetic particle spectroscopy, magnetic coupling, agglomeration, aggregation

## Abstract

Tracer and thus signal stability is crucial for an accurate diagnosis via magnetic particle imaging (MPI). However, MPI-tracer nanoparticles frequently agglomerate during their in vivo applications leading to particle interactions. Here, we investigate the influence of such magnetic coupling phenomena on the MPI signal. We prepared and characterized Zn_0.4_Fe_2.6_O_4_ nanoparticles and controlled their interparticle distance by variying SiO_2_ coating thickness. The silica shell affected the magnetic properties indicating stronger particle interactions for a smaller interparticle distance. The SiO_2_-coated Zn_0.4_Fe_2.6_O_4_ outperformed the bare sample in magnetic particle spectroscopy (MPS) in terms of signal/noise, however, the shell thickness itself only weakly influenced the MPS signal. To investigate the importance of magnetic coupling effects in more detail, we benchmarked the MPS signal of the bare and SiO_2_-coated Zn-ferrites against commercially available PVP-coated Fe_3_O_4_ nanoparticles in water and PBS. PBS is known to destabilize nanoparticles mimicking an agglomeration in vivo. The bare and coated Zn-ferrites showed excellent signal stability, despite their agglomeration in PBS. We attribute this to their aggregated morphology formed during their flame-synthesis. On the other hand, the MPS signal of commercial PVP-coated Fe_3_O_4_ strongly decreased in PBS compared to water, indicating strongly changed particle interactions. The relevance of this effect was further investigated in a mammalian cell model. For PVP-coated Fe_3_O_4_, we could detect a strong discrepancy between the particle concentration obtained from the MPS signal and the actual concentration determined via ICP-MS. The same trend was observed during their MPI analysis; while SiO_2_-coated Zn-ferrites could be precisely located in water and PBS, PVP-coated Fe_3_O_4_ could not be detected in PBS at all. This drastically limits the sensitivity and also general applicability of MPI using such standard commercial tracers and highlights the advantages of our flame-made Zn-ferrites concerning signal stability and ultimately diagnostic accuracy.

## Introduction

Magnetic particle imaging (MPI) is an emerging diagnostic method with various promising application areas. It is based on the non-linear response of magnetic nanoparticles to an applied oscillating magnetic field.^[1]^ The tracer particles are injected and accumulate via active or passive targeting mechanisms at the site of interest. A static magnetic field with strong gradients is applied to saturate the majority of the particles in the tissue and only creating a small field-free region.^[2]^ An oscillating magnetic field is superimposed to the static magnetic field and only the signal of the MPI tracers in the field-free region is detected via coils. The field-free region is then swept over the entire volume of interest (field of view) to obtain a three-dimensional image.^[3]^ The main advantage of MPI lies in its high sensitivity down to 10^-6^ – 10^-8^ M.^[4]^ As only the tracers are imaged, there is no disturbance through background signals created by the tissue itself. MPI does not require harmful ionizing radiation, in contrast to frequently used positron emission tomography (PET). Furthermore, this comparably inexpensive imaging technique stands out due to its high spatial and temporal resolution outperforming clinically established methods such as magnetic resonance imaging (MRI) or computer tomography (CT).

The overall MPI performance strongly depends on the exact tracer characteristics.^[4]^ The magnetization of these nanoparticles is frequently described via the Langevin function.^[2]^ However, the latter neglects the relaxation mechanisms (i.e. Néel- and Brownian relaxation) present in larger particles and caused by thermal perturbations. These result in a delay of the magnetic response to the field, which drastically complicates the tracer dynamics. Surprisingly, research on the practical optimization of these nanoparticles is still in its infancy. Previous studies have frequently used commercially available materials or clinically approved MRI contrast agents based on iron oxides. While this approach would potentially allow a swift translation of the tracers into clinics provided that human-sized MPI scanner become feasible, the investigated systems are far from ideal for this application. In fact, according to simulations, only 3% of the iron mass of the clinically approved Ferucarbotran (Resovist) dispersion contribute to the MPI signal. The major part was suggested to remain inactive due to its too small size.^[1]^ Follow-up studies indicated that this mass might increase by a factor of 30 due to particle interactions but remains inefficiently low.^[5]^ While ferucarbotran based nanoparticles are still used as the standard MPI tracers, attempts to improve signal strength through material engineering have been reported. So far, they have focused on size-optimizations of polymer-coated iron oxide nanoparticles.^[6]^ However, a deeper understanding of the involved mechanisms especially under physiologically relevant conditions is yet to be achieved.

Next to signal strength, also, signal stability is crucial for a reliable and accurate diagnosis via nanoparticle-based imaging methods. In this regard, particle interactions play a fundamental role.^[7]^ Nanoparticles are frequently characterized as monodisperse non-interacting spheres in aqueous dispersions ex vivo. Upon their injection in vivo or their exposure to cells in vitro^[8,9]^, they agglomerate due to the harsher conditions faced.^[10]^ These changes in the surroundings include variations in the salt concentration, pH, or protein adsorption. As a result of the agglomeration, the primary particles are in close contact and start interacting. Magnetic nanoparticles will couple, which strongly alters their overall magnetic properties.^[11]^ This phenomenon has been investigated for various sensing applications via MRI^[12,13]^ or MPS^[14]^. Interestingly, however, it has been frequently neglected otherwise. Polymer coatings only slightly improve the stability and in long run have been shown to even disintegrate exposing the bare particle surface.^[15]^

We have investigated the importance of interactions between magnetic particles used as MRI contrast agents.^[16]^ Iron oxide nanoparticles with a size below approximately 5 nm become paramagnetic and are thus excellent T1 MRI contrast agents. We showed, however, that bare iron oxide particles in this size regime strongly couple, which increases their magnetic size and renders them superparamagnetic/ferromagnetic instead of paramagnetic. We demonstrated that introducing a silica matrix as a spacing material can effectively counteract particle coupling. Through this, we sterically hindered the magnetic cores to interact and allowed the system to act as a high-performing contrast agent.

Similar studies for MPI tracers are scarce. Recently, such magnetic interactions have been termed superferromagnetism and investigated for signal enhancements.^[17]^ Moreover, Khandhar et al. showed that the signal of commercial polymer-coated nanoparticles decreases by 53% in cell-culture medium compared to water.^[18]^ They attributed this reduction to particle interactions and resulting changes in magnetic relaxation.^[19]^ The obtained signal can no longer be assigned to a specific particle concentration via a calibration curve, as the latter strongly depends on the exact particle state, which is typically unknown. This was briefly investigated by encapsulating commercial polymer coated iron oxide particles into red blood cells.^[20]^ The obtained MPS signals of the cells with the tracers revealed a strong discrepancy to the reference samples in water. Such differences between effective and predicted tracer concentrations are highly problematic. It drastically increases the risks of false diagnoses caused by over- or underpredictions of tracer quantities.

In this work, we aim to investigate and mitigate this risk by controlling particle interactions through core-shell structures. To this end, we utilize the versatile flame-synthesis, which has been previously investigated for the synthesis of a wide range of magnetic nanomaterials of different sizes^[21]^, compositions^[22]^, and morphologies^[23]^. More specifically, we investigate SiO_2_-coated non-stoichiometric Zn-ferrites with improved magnetic properties compared to pure iron oxides and high biocompatibility.^[24]^ We focus on analyzing and optimizing MPS signal stability under physiologically relevant conditions.

## Experimental methods

### Particle synthesis

Zn_0.4_Fe_2.6_O_4_ nanoparticles were produced by flame spray pyrolysis, as previously described.^[23,24]^ A liquid precursor solution was fed at 5 mL/min through a capillary and dispersed by 5 L/min O_2_ into fine droplets. These are ignited by flaming CH_4_/O_2_ mixture (1.5 and 3.2 L/min, respectively), which then leads to the primary particle formation. The flame is further surrounded by 40 L/min of O_2_ sheath gas. The particle stream is guided via a quartz glass tube (length 20 cm, inner diameter 45 mm) through the SiO_2_ coating ring. The latter has 16 openings facing 30° downwards the particle stream. HMDSO (hexamethyldisiloxane, Sigma-Aldrich) vapor is injected through these openings, which condenses on the freshly formed core particles and forms the SiO_2_ layer. The HMDSO vapor is fed via an N_2_ stream saturated in a bubbler and further diluted by 16 L/min N_2_. The SiO_2_ content in the final product is controlled through the N_2_ bubbler flow and calculated at saturation conditions (T = 20 °C, p_vapor,HMDSO_ = 43 mbar, wt% SiO_2_ = m_SiO2_ / (m_SiO2_ + m_Zn0.4Fe2.6O4_). The stream of the now SiO_2_ coated Zn_0.4_Fe_2.6_O_4_ particles is guided via another quartz glass tube (length 30 cm, inner diameter 45 mm) onto a glass fiber filter (Whatman GF6, 257 mm diameter), where the particles are collected with the help of a vacuum pump (Busch, Seco SV 1040C).

Before the synthesis, the precursor solution for the Zn_0.4_Fe_2.6_O_4_ nanoparticles was prepared at a concentration of 0.2 mol_Zn+Fe_ / L in a 3:1 volume ratio of xylene and acetonitrile (both Sigma-Aldrich). Iron acetylacetonate (Sigma-Aldrich) and zinc acetylacetonate (Sigma-Aldrich) were dissolved accordingly via magnetic stirring for 1 h at room temperature. HMDSO was utilized in the bubbler as supplied.

PVP-coated Fe_3_O_4_ was purchased from Nanocomposix (NanoXact Magnetite Nanoparticles – PVP −20 mg/mL in aqueous 2 mM sodium citrate, 20 nm ± 5 nm).

### Characterization

X-ray diffraction patterns were recorded on a Bruker D2 Phaser (30 kV, mA). The obtained data was analyzed via Diffrac Eva and TOPAS 4.2. software. XRD intensity spectra were fitted over all angles (2θ) to the standard pattern of magnetite (Fe_3_O_4_, IDSC: 84611) and crystal size computed via Rietveld refinement.

Scanning transmission electron microscopy (STEM) was performed on a Talos F200x microscope (ThermoFisher, field emission gun at 200 kV) with four attached silicon drift detectors. Samples were prepared by dispersing nanoparticles in mili-Q ultra pure water. A dispersion drop was deposited on a carbon-coated copper grid (EMR, Lacey Carbon Film 200 Mesh Copper) and then carefully dried. The primary core particle size distribution was estimated by measuring the longest axis of 176 bare and 103 70 wt% SiO_2_ coated Zn_0.4_Fe_2.6_O_4_ nanoparticles using ImageJ software and convergence of geometrical standard deviation was ensured. Log-normal size distribution was assumed.

Dynamic light scattering analysis for hydrodynamic size and ζ-potential was conducted on a Zetasizer (Nano ZS90 Malvern Instruments). Samples were prepared by dispersing the nanoparticles in mili-Q ultra-pure water or PBS at a concentration of 0.1 mg/mL via 10 minutes of sonication (Ultrasonic Processor, pulses: 28s/2s, 90% Amplitude).

Magnetic properties were assessed via VSM on a Quantum Design Physical Property Measurement System. The samples were prepared by measuring the weight of the particles and placing them between two capillaries. The magnetization versus applied field curves (MH-curves) was assessed at 300 K. The applied field ranged from 3000 mT to −3000 mT. Zero-field cooled (ZFC) and field cooled (FC) curves were obtained by varying the temperature in the range from 10 K to 300 K and applying a magnetic field of 150 Oe for FC.

### Magnetic particle spectroscopy and imaging

To investigate the particle response under a sinusoidal magnetic field, a custom-made and calibrated magnetic particle spectrometer (MPS) similar to a previous study^[25]^ was used. An MPS can be interpreted as an MPI-scanner without a spatial encoding. A time-dependent magnetization curve is measured via coils by exciting the particles in one dimension. The base frequency of the magnetic field was set to 26.042 kHz and the amplitude to 20 mT. For further understanding of the particle behavior the spectrum of the magnetization curve is investigated in the frequency domain. The harmonics of the base frequency can be used to compare the response of different particles and their general suitability for MPI.

In addition to the particle characterization via onedimensional MPS exciation, the particles were investigated regarding their applicability for imaging using a preclinical MPI-scanner (Bruker, Germany). To this end two-dimensional system matrices were measured using a point source which is driven through the field of view by a roboter. A system matrix describes the systems answer of every discrete voxel for every frequency component and is needed to reconstruct the particle signal from the received voltage signal. The system matrices of 8 μL cubic particle samples (17 g/L in water) were measured using a gradient of 2 T/m and a drive field strength of 12 mT leading to a field of view of 24 x 24 mm discretized in 13 x 13 voxels of size 1.85 x 1.85 mm. A signal-to-noise threshold of 5 was applied together with a minimum frequency of 80 kHz. For a good spatial resolution of the reconstructed image, particles should give clear system matrix patterns for as much frequency components as possible. The investigation of the decay of the system matrices amplitude over the harmonics is therefore a crucial measure to compare different particles in their suitability for MPI imaging. Images were reconstructed with the open-access Julia package MPIReco^[26]^ using an iterative regularized Kaczmarz algorithm, a relaxation parameter λ of 0.001 and 100 iterations.

For the in vitro experiments, cells were seeded at 250’000 per T25 flask in 7.2 mL medium and left to attach for 24 h. 800 μL of 1 mg/mL nanoparticle dispersion in ultra-pure water (after ultra-sonication) were added for a final concentration of 0.1 mg/mL. After 24 h of incubation cells were washed twice with PBS and then trypsanized using 500 μL Trypsin. Trypsinization was then stoped with 1 mL MEM. Thereafter, the liquids were removed and replaced 500 μL 4%PFA. After 4 h PFA was replaced with 1.5 mL PBS. The cells were spinned down and transfered in 100 μL to fresh Eppendorf tubes. Next, the cells were centrifuged and the cell pellet was resuspended in 20 μL of 0.5 % Agar (diluted from hot 1.5%Agar using prewarmed PBS) and the samples were put in a freezer for 10 min. Finally, 100 μL Mowiol was added on top for protection and left to harden over night at room temperature.

For ICP-OES samples, all washings as well as the cell pellets were collected seperately to distinguish intra- and extracellular particles and digested at 250 °C in concentrated nitric acid and hydrogen peroxide using a microwave (turboWAVE Inert, MLS GmbH, Leutkirch, Germany). Samples were diluted using ultrapure water to reach a final concentration of 2% nitric acid. Fe and Zn contents were determined by an ICP-OES 5110 (Agilent, Basel, Switzerland) instrument with external calibration ranging from 0 to 5 ppm.

## Results and discussion

To investigate the importance of particle interactions on MPI, bare and SiO_2_ coated Zn_0.4_Fe_2.6_O_4_ nanoparticles were prepared by scalable flame synthesis. The SiO_2_ shell is applied in situ^[27]^, which allows the coating of individual core particles and thus gives control over their interparticle (core-to-core) distance. Figure 1 (top) shows scanning electron microscopy images of the prepared bare (a) and 70 wt% SiO_2_-coated (b) Zn_0.4_Fe_2.6_O_4_ nanoparticles. The homogenous SiO_2_ shell surrounding the predominantly hexagonal particles is visible. This leads to a clear separation of the magnetic Zn_0.4_Fe_2.6_O_4_ core in comparison to the bare sample. Both images show that the primary particles are connected via sinter necks to larger aggregates, which is characteristic for flame-made nanomaterials^[28]^.

**Figure 1.**
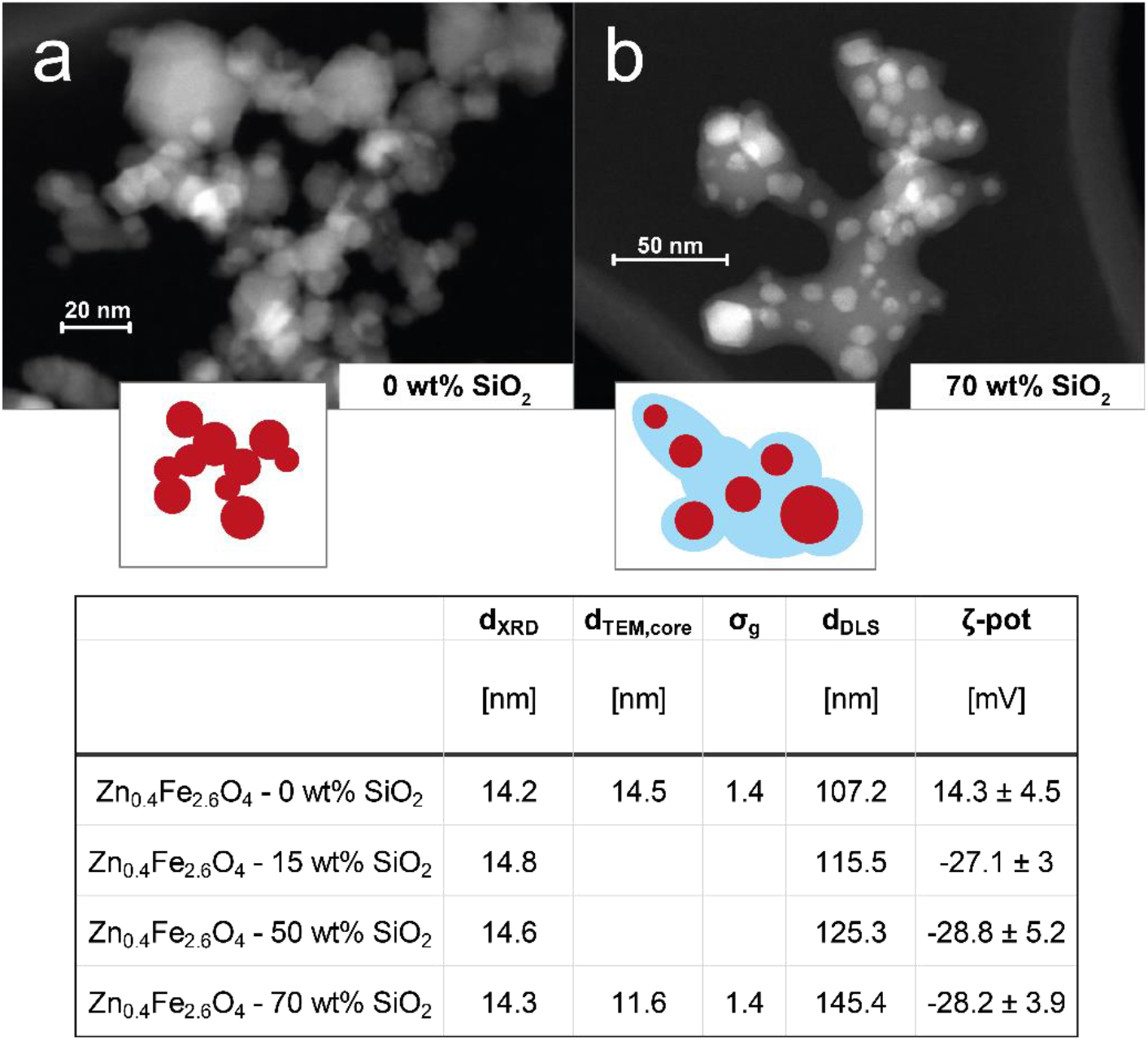
STEM images of as-prepared (a) bare and (b) 70 wt% SiO_2_-coated Zn_0.4_Fe_2.6_O_4_ nanoparticles. Insets schematically depict characteristic morphology. Table (below) summarizes morphological characteristics of the different prepared particles: crystal sizes (d_XRD_), geometric mean primary particle sizes of core (d_TEM,core_), geometrical standard deviations of primary particle sizes of core (σ_g_), geometric means of hydrodynamic diameter (d_DLS_), ζ-potentials (ζ-pot).

The as-prepared nanoparticles were further analyzed concerning their morphology and composition. The table in Figure 1 (bottom) summarizes the results. The XRD patterns (see Figure S1) of the bare and all SiO_2_-coated particles indicate the desired spinel ferrite structure^[29]^. The crystal size lies between 14 and 15 nm is unaffected by the SiO_2_ coating. This is in agreement with previous reports on in situ SiO_2_ coated flame-made nanoparticles^[30]^ and shows that the coating was applied after the core particle formation had been completed. The geometrical mean core diameters determined from TEM analysis (d_TEM,core_) are in agreement with the crystal sizes and show a geometric standard deviation σ_g_ of 1.4. The agglomerate size measured in water (d_DLS_) slightly increases for increasing SiO_2_ contents, as expected for a thicker shell. The complete coating even for the lowest SiO_2_ content is suggested by the shown constant ζ-potentials for all coated samples. The ζ-potential values show a distinct decrease from +14 mV for bare Zn_0.4_Fe_2.6_O_4_ to −28 mV for all SiO_2_-coated particles, in agreement with the literature.^[24]^

Next, the prepared nanomaterials were analyzed in detail concerning their magnetic properties. Figure 2a shows the magnetization curves normalized to the total sample mass (incl. SiO_2_). All samples show an S-shaped data set characteristic to ferro- and ferrimagnetic systems. There is no major hysteresis detectable indicating that the magnetic sizes of the particles are within the superparamagnetic range (i.e. < appr. 25 nm for Fe_3_O_4_^[31]^). As summarized in Table 1, the coercivities H_c_ decrease from 0.86 mT for the bare to 0.39 mT for the 70 wt% SiO_2_-coated sample. While the values for the saturation magnetization M_s_ per sample mass decrease for increasing SiO_2_ contents as expected, they remain constant at approximately 60 emu/g if normalized to the nominal core mass. This value corresponds to the literature^[24]^ and suggests that the volume and the composition of the magnetic core are not affected by the SiO_2_ coating, which is in agreement with the constant d_XRD_.

**Figure 2.**
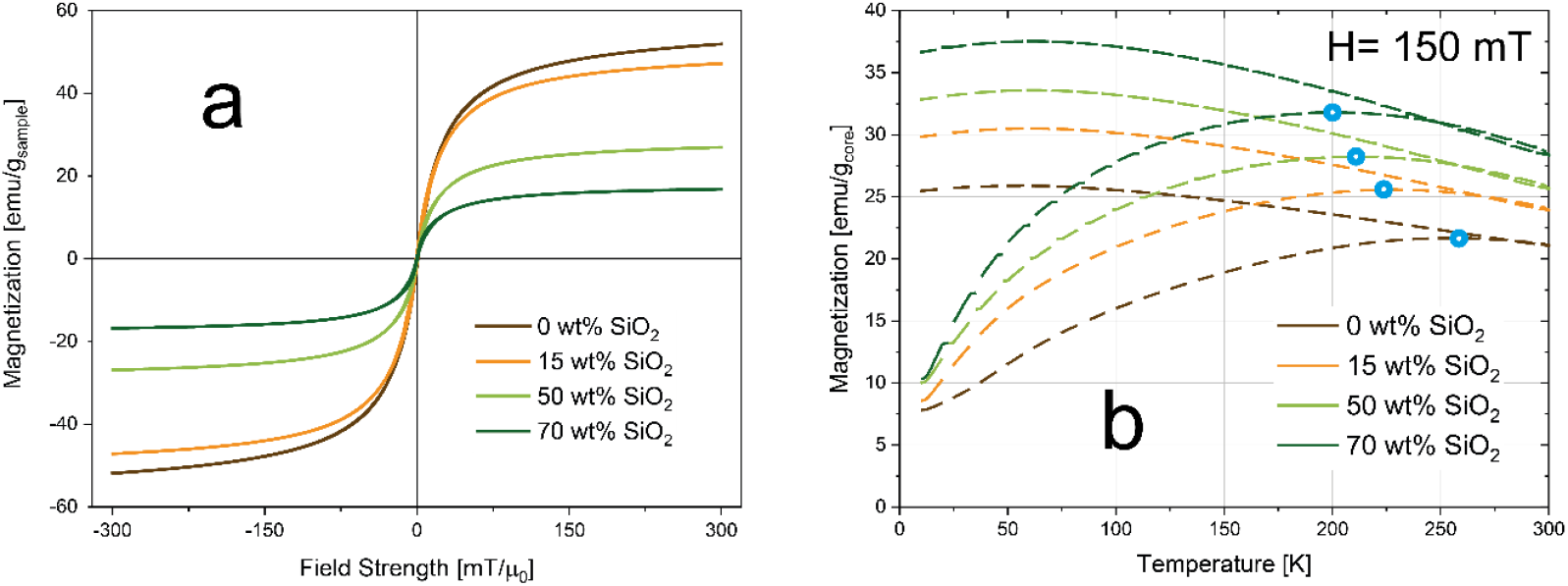
(a) Magnetizations per mass of overall sample as function of applied field. (b) Magnetizations per mass of overall sample as a function of temperature at a constant applied field of 150 mT (i.e. field-cooling curves). Blocking temperatures are indicated by a blue hollow sphere.

**Table 1.** Summary of magnetic properties extracted from magnetization and field-cooling curves: saturation magnetizations (M_s_) per mass of overall samples and per nominal mass of magnetic core, coercivities (H_c_), magnetic susceptibilities (χ), blocking temperatures (T_B_).

| | $M_s$ | | $H_c$ | $\chi$ | $T_B$ |
| --- | --- | --- | --- | --- | --- |
|  | [emu/g<sub>sample</sub>] | [emu/g<sub>core</sub>] | [mT] | [1/mT] | [K] |
| Zn<sub>0.4</sub>Fe<sub>2.6</sub>O<sub>4</sub> - 0 wt% SiO<sub>2</sub> | 59 | 59 | 0.86 | 0.03 | 252 |
| Zn<sub>0.4</sub>Fe<sub>2.6</sub>O<sub>4</sub> - 15 wt% SiO<sub>2</sub> | 52.7 | 62 | 0.72 | 0.036 | 227 |
| Zn<sub>0.4</sub>Fe<sub>2.6</sub>O<sub>4</sub> - 50 wt% SiO<sub>2</sub> | 29.7 | 59.4 | 0.31 | 0.047 | 215 |
| Zn<sub>0.4</sub>Fe<sub>2.6</sub>O<sub>4</sub> - 70 wt% SiO<sub>2</sub> | 18.1 | 60.3 | 0.39 | 0.059 | 201 |

The susceptibility χ increases for thicker SiO_2_ shells. This trend is also evident in Figure 2b, which shows the field-cooled curves i.e. the magnetization normalized by the nominal core mass at an applied field of 150 mT as a function of temperature for all samples. The magnetization at 300 K increases similarly to χ, indicating a more pronounced magnetic response for the coated samples at low applied fields compared to the bare particles. The blocking temperatures (T_B_, blue marks) refer to the transition from the magnetically blocked to the superparamagnetic state. At this temperature, the magnetic dipole moments possess enough thermal energy to overcome the applied field and can thus rotate freely. The T_B_ values shift towards lower temperatures for higher SiO_2_ contents. This indicates reduced magnetic dipole interactions due to increased interparticle distances caused by the coating.^[32]^

The above-discussed influence of particle interactions on magnetic properties has been measured under quasi-static applied fields via vibrating sample magnetometry (VSM). However, MPI is utilizing oscillating magnetic fields at drastically higher frequencies, which strongly affects the magnetic response due to the involved dipole relaxation mechanisms. To investigate this influence in detail, micromagnetic simulations were performed via the Object Orientated Micro Magnetic Framework (OOMMF) software (Figure S2).^[33]^ The results also indicate a strong dependence of the coercivity on the exact inter-particle distance. This, however, corresponds only partly to the experimentally obtained VSM and MPS data presented in this work.

The prepared particles were then analyzed concerning the imaging performance via magnetic particle spectroscopy (MPS, i.e. MPI without spatial encoding). Figure 3 summarizes the MPS performance of the prepared bare and SiO_2_-coated Zn_0.4_Fe_2.6_O_4_. All measurements were performed as aqueous dispersion at a constant nominal core concentration to directly compare the magnetic properties. Figure 3a shows the obtained voltage signal over time and Figure 3b the absolute signal as a function of the applied field. The signal amplitude increases clearly with increasing SiO_2_ shell thickness, despite the constant amount of magnetic mass. Although measured at different frequencies this is in excellent agreement with the susceptibility trend shown above (Table 1). The magnetic spectrum shown in Figure 3c confirms the observations by showing clear differences between the bare and all SiO_2_-coated samples over all frequencies. However, the influence of the coating thickness is only minor. Figure 3d depicts the hysteresis curves derived from the MPS signals. The maximal magnetizations remain almost constant for all samples. Differences are mostly detectable in the slopes as also shown in Figure 3b. Interestingly, coercivity values remain almost constant for all samples. This is in strong contrast to the obtained computational results shown in Figure S2. Especially, the similar value for the bare sample compared to all SiO_2_-coated particles cannot be explained.

**Figure 3.**
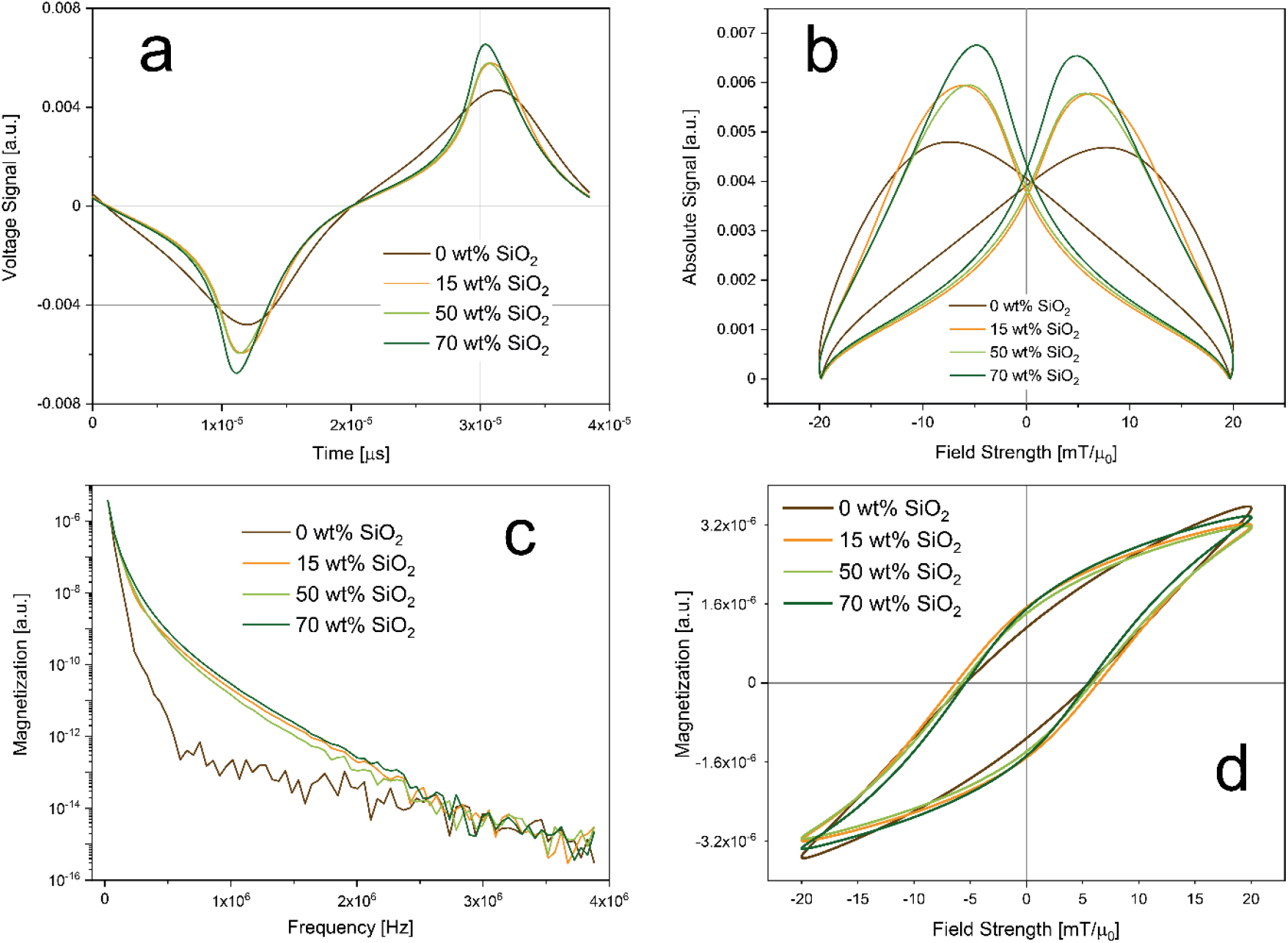
MPS results: (a) measured signal as function of time; (b) measured signal as a function of applied field (i.e. point spread function); (c) magnetization as a function of frequency (i.e. magnetic spectrum); (d) magnetization as a function of applied field strength (i.e. hysteresis). Measurements were conducted at 20 mT with a frequency of 26 kHz.

To investigate this behavior and the effect of magnetic interactions in more detail, we conducted MPS measurements in different dispersion media. The obtained signal harmonics show good linear agreement with the particle concentration down to 16 μg/mL. Even though frequently used in literature, the voltage amplitude is sufficiently linear only down to approximately 1 mg/mL (see Figure S3). Figure 4a depicts the 3^rd^ harmonic MPS signals as a function of particle concentration in water and PBS. Data is shown for bare and SiO_2_-coated (70 wt%) Zn_0.4_Fe_2.6_O_4_, as well as commercial PVP-coated Fe_3_O_4_. The latter belongs to the group of commonly used MPI tracers in literature (i.e. polymer-coated iron oxides).^[4]^Although, PVP-coated Fe_3_O_4_ has the strongest signal in water it also loses most of its performance in PBS. Our flame-made bare and SiO_2_-coated Zn-ferrites, on the other hand, show comparably constant calibration lines. In the 15^th^ harmonic (Figure 4b), the SiO_2_-coated Zn-ferrite particles distinctively outperform the other systems while still attaining good stability over the dispersion medium. This trend is further analyzed in Figure 4c, which depicts the slopes of the calibration lines as a function of the signal harmonics for all samples in water and PBS. PVP-coated Fe_3_O_4_ nanoparticles show steeper slopes at lower harmonics than the flame-made particles. For the 7^th^ and higher harmonics, however, SiO_2_-coated Zn_0.4_Fe_2.6_O_4_ nanoparticles show the best performance. The signal of commercial polymer coated particles continuously decreases reaching a signal value of 10^-12^ at the 13^th^ harmonic which is typically used as the detection limit. At this point, SiO_2_-coated Zn-ferrite nanoparticles outperforms PVP-coated Fe_3_O_4_ by more than two orders of magnitude. The signal stability over the dispersion medium is further analyzed in Figure 4d, which shows the ratio of the calibration line slops in PBS over water. Our flame-made bare and SiO_2_-coated Zn-ferrites show excellent signal stability throughout all harmonics. Interestingly, the bare particles attain slightly steeper calibration curver in PBS compared to H_2_O, which presently cannot be explained. On the other hand, PVP-coated Fe_3_O_4_ suffers from a big discrepancy between the signals measured in H_2_O and PBS. The cross-over at the 13^th^ harmonic most likely is an artifact from the weak signal (< 10^-12^) at these frequencies.

**Figure 4.**
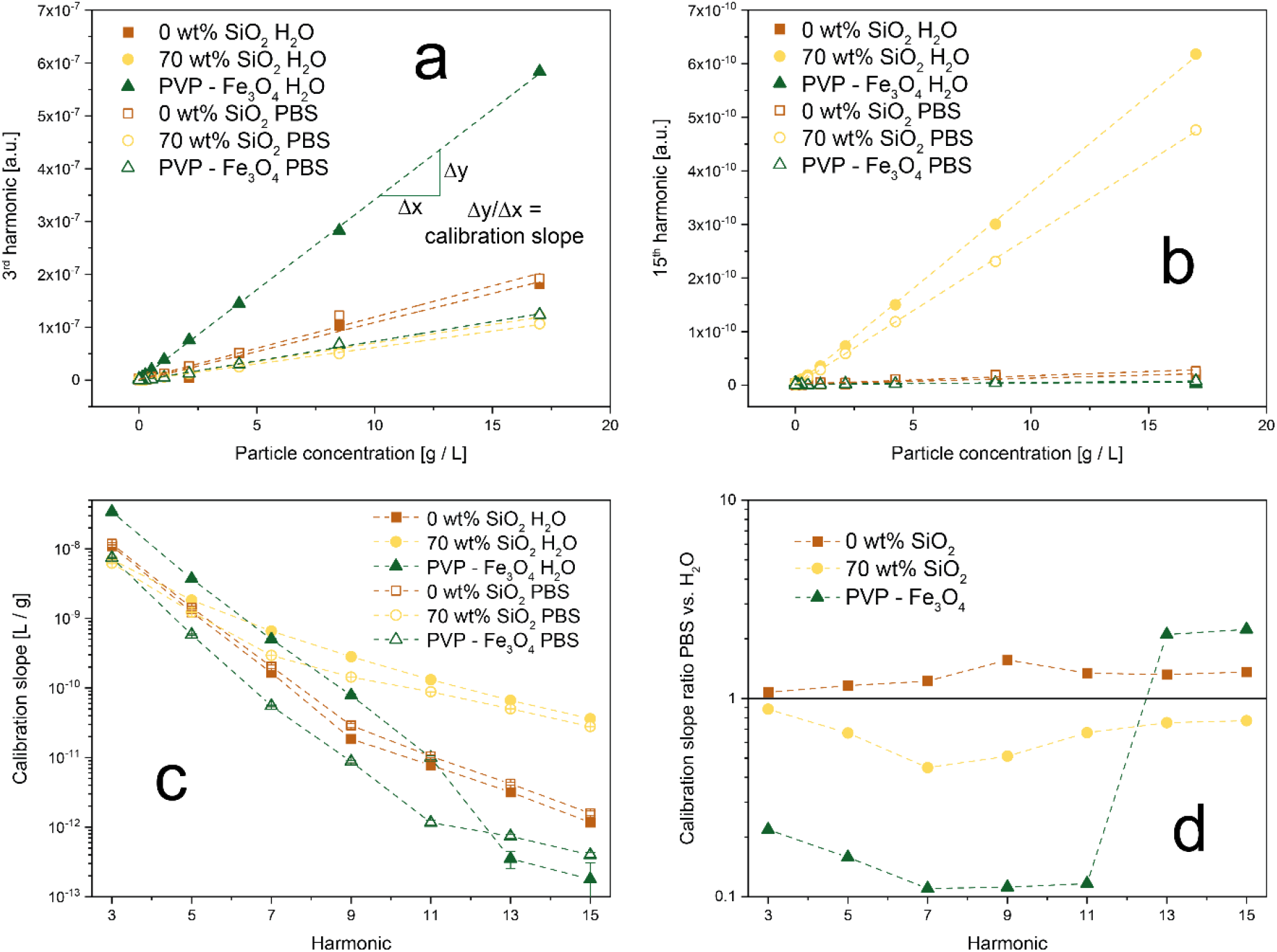
Comparison of signal stability as a function of dispersion medium for synthesized Zn-ferrites and commercial magnetite nanoparticles. Calibration curves for (a) 3^rd^ and (b) 15^th^ harmonic for bare and SiO_2_-coated (70 wt%) Zn_0.4_Fe_2.6_O_4_, as well as commercial PVP-coated Fe_3_O_4_ dispersed in H_2_O (closed symbols) or PBS (open symbols). (c) The slope of the calibration lines as a function of the harmonics for all samples in H_2_O and PBS. (d) The ratio of the calibration slopes in PBS over H_2_O for all particles. A value close to 1 indicates a stable signal.

The aformentioned results can be explained by changes in their state of agglomeration. Khandhar et al. reported a similar trend with MPS signal voltage decreases of 8%, 53%, and 74% in blood, cell-culture medium, and 1 wt% agar gel, respectively.^[18]^ Figure S3 shows the hydrodynamic diameters of the samples measured in water and PBS. While the flame-made particles attain a size of approximately 100 nm in H_2_O, the commercial PVP-coated Fe_3_O_4_ has a size of approximately 30 nm. Interestingly, upon exposure to PBS bare Zn-ferrite and the commercial PVP-coated Fe_3_O_4_ show a strong size increase to about 1000 nm. This increase corresponds to literature and can be explained by the high ionic strength of PBS and the thus weakened repulsive forces leading to stronger agglomeration.^[34]^ The hydrodynamic diameter of SiO_2_-coated (70 wt%) Zn_0.4_Fe_2.6_O_4_ nanoparticles, on the other hand, decreases. Although the increased agglomeration is observed also for bare Zn-ferrites, instabilities in the MPS signal are only observable for PVP-coated Fe_3_O_4_. Firstly, we attribute this to a stronger size increase of the commercial particles in PBS compared to water. Secondly, this can be explained by morphological differences between materials. The prepared bare and SiO_2_-coated Zn-ferrites show a strongly aggregated structure (Figure 1) characteristic of flame-made nanoparticles.^[28]^ This is a result of their high-temperature synthesis leading to the formation of strong sinter-necks between the primary particles, which remain stable even during harsh dispersion conditions. As a result, the magnetic cores have a fixed distance (bare: touching, SiO_2_-coated: separated) and thus magnetic coupling originating from their synthesis. Additional agglomeration in dispersion does not substantially affect these interactions. PVP-coated Fe_3_O_4_, on the other hand, is characterized as monodisperse non-interacting particles in water. Upon their exposure to PBS, they strongly agglomerate leading to particle interactions despite the polymer coating. Ultimately, this results in a highly unstable MPS signal. This experiment simplistically mimics the conditions faced by nanoparticles in vitro and in vivo, causing unavoidable agglomeration.^[7]^ This is especially problematic as the obtained MPI signals are typically assigned to particle concentrations via external calibration curves.

To analyze the limitations of quantitative MPS/MPI under more realistic conditions, bare and SiO_2_-coated Zn-ferrites, as well as commercial PVP-coated Fe_3_O_4_ nanoparticles were incubated with cells for 24 h. The precise amount of particles taken up by the cells was determined via ICP-MS. The cell samples were then analyzed via MPS. Signal harmonics or particle concentrations predicted through the calibration lines were then compared to the effective values determined by ICP-OES. Figure 5a shows the ratio of the MPS signal harmonics determined through the calibrations using the exact particle concentration over the actual MPS signal. Throughout all relevant harmonics, SiO_2_-coated Zn-ferrites show the best accuracy, with comparable signal ratios for H_2_O and PBS dispersions. The good performance of PVP-coated Fe_3_O_4_ at higher harmonics can be explained again by low signal intensities. A similar trend can be observed in Figure 5b, which shows the error of the particle concentrations determined via the MPS signal. SiO_2_-coated Zn-ferrites show a comparably good prediction of the particle concentration throughout all signal harmonics. PVP-coated Fe_3_O_4_, on the other hand, drastically underpredicts the concentration by at least 85%. At higher harmonics and for all calibration curves in H_2_O these particles perform especially poorly with errors above 100% (data not shown).

**Figure 5.**
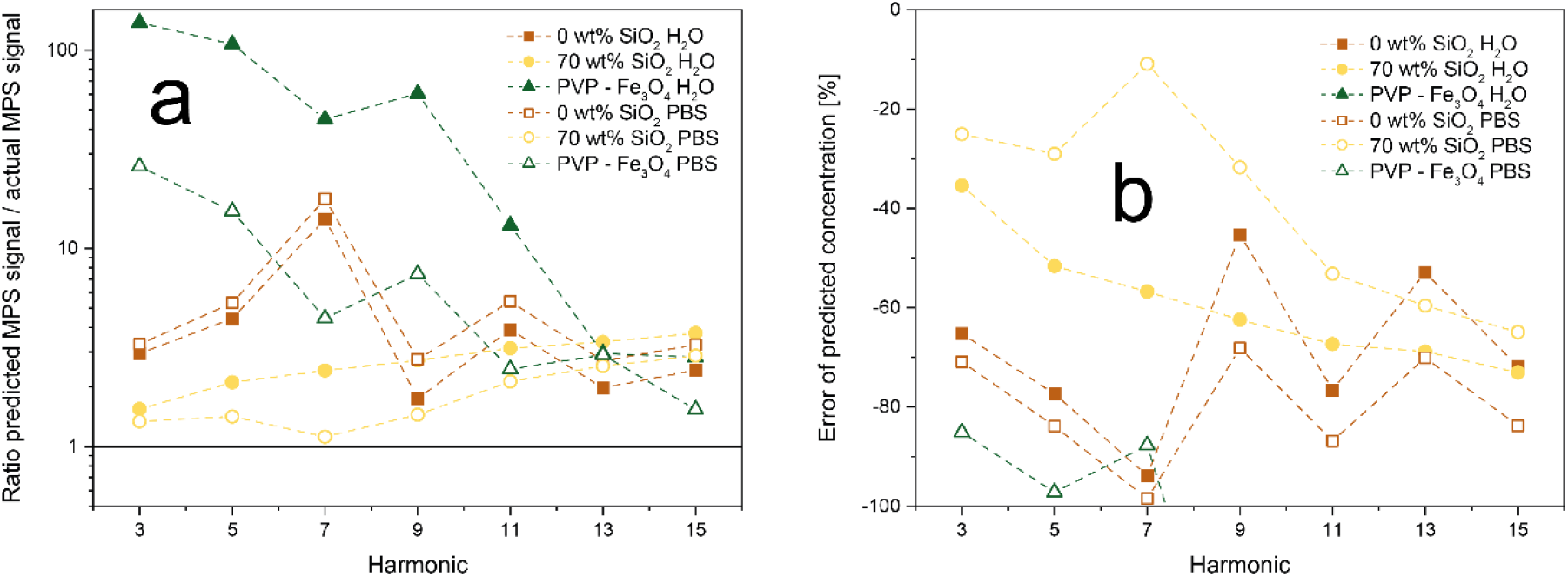
Signal stability in vitro. Particles were incubated with cells for 24h and their concentration and MPS signal after washing were determined. Data is shown for bare and SiO_2_-coated (70 wt%) Zn_0.4_Fe_2.6_O_4_, as well as commercial PVP-coated Fe_3_O_4_ dispersed in H_2_O (closed symbols) or PBS (open symbols). (a) Ratio of MPS signals predicted at the measured particle concentration through calibration curves and actual measured MPS signal. (b) Error of particle concentration determined via MPS calibration curves.

However, the underprediction of the concentrations can be observed for all samples and both water and PBS calibrations curves. This can be explained by the fixation of the cells and thus also the particles in a polymer, which suppresses Brownian relaxation mechanisms and in the following the overall MPS signal. For SiO_2_-coated Zn-ferrite, the PBS calibrations result in better accuracy. Most likely, this is a result of the increased particle agglomeration in PBS better mimicking the in vitro morphology. Nevertheless, it has to be mentioned that even for the best performing SiO_2_-coated Zn-ferrites in PBS the average error in concentration over all harmonics lies at around 40%. This value increases to 60% when using the calibration in water. This highlights the importance of magnetic particle interaction and shows the current limitations of quantitative MPS and MPI.

In a final step, we investigate the consequences of the particle and thus signal stability on the actual magnetic particle imaging performance. To this end, we obtained the system matrix at 13^2^ different locations in a 2D MPI setup for SiO_2_-coated (70 wt%) Zn_0.4_Fe_2.6_O_4_, as well as commercial PVP-coated Fe_3_O_4_ dispersed in H_2_O or PBS. Bare Zn-ferrites could not be analyzed due to too strong sedimentation of particles during the measurement time. Figure 6a shows the obtained system matrices for the 66^th^ frequency component (k) for all samples. Distinct patterns can be observed for the SiO_2_-coated Zn-ferrites indicating a good signal-to-noise ratio both in water and PBS. In contrast, the polymer-coated particles show the characteristic system matrix features only in aqueous dispersion, while no signal could be detected in PBS. These observations are confirmed by Figure 6b, which shows the MPI signal as a function of the harmonics for all samples. PVP-coated Fe_3_O_4_ in H_2_O results in a high signal at low harmonics, which quickly declines thereafter. Flame-made SiO_2_-coated Zn-ferrites show excellent signal strength in both media especially at higher harmonics, where they outperform the commercial polymer-coated particles. Differences between the signals in PBS versus water are depicted in Figure 6d. SiO_2_-coated (70 wt%) Zn_0.4_Fe_2.6_O_4_ particles show excellent signal stability over the entire harmonic range. They distinctively outperform PVP-coated Fe_3_O_4_, which shows signal differences of up to two orders of magnitude. This discrepancy is further reflected in the reconstructed images depicted in Figure 6c. As expected, SiO_2_-coated Zn-ferrites can be precisely located in water and PBS, while only aqueous PVP-coated Fe_3_O_4_ results in a satisfactory image.

**Figure 6.**
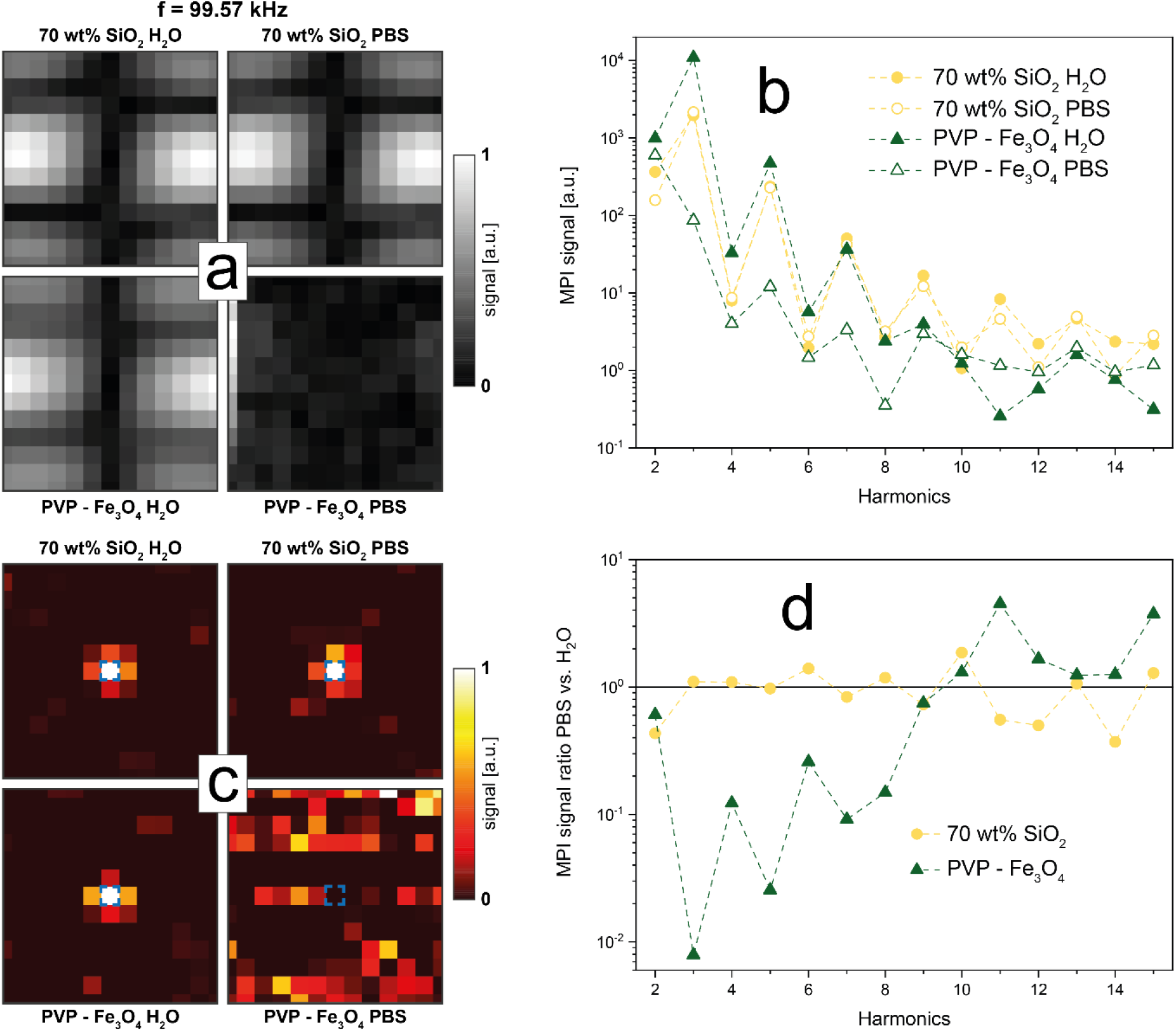
Influence of dispersion medium on MPI quality of SiO_2_-coated (70 wt%) Zn_0.4_Fe_2.6_O_4_, as well as commercial PVP-coated Fe_3_O_4_ dispersed in H_2_O or PBS at 17 g / L. (a) Absolute values of the system matrices at 99.57 kHz. Values are normalized to sample maximum. (b) Signal at the particle location as a function of the signal harmonics. (c) Reconstructed images of all samples. Signals are normalized to sample maximum. Blue square indicates particle location. (d) Ratio of signal measured in H_2_O over PBS at the particle location as a function of the signal harmonics.

## Conclusions

Nanoparticle agglomerations as well as the resulting particle interactions in vitro and in vivo are frequently neglected. However, their consequences ranging from changes in bio-distribution or altered magnetic characteristics, as described here, are severe. Here, we report that magnetic particle imaging is specifically affected by such magnetic coupling effects.

We found a distinctly better MPS signal for SiO_2_-coated compared to bare Zn-ferrites. However, the exact thickness of the shell did not substantially influence the signal. Most importantly, both here prepared systems showed a constant MPS signal, irrespective of their dispersion medium. Conversely, frequently used polymer-coated Fe_3_O_4_ nanoparticles drastically lost signal intensity at the same particle concentration in PBS compared to water. We explain this by the pre-aggregated state and the strongly changed particle interaction of our Zn-ferrites and the commercial particles, respectively.

Signal stability is crucial for the exact quantification of tracer amounts during the magnetic particle imaging process. Latter is a frequently mentioned key advantage of MPI compared to other imaging methods, which, however, might be limited when using gold-standard polymer-coated nanoparticles. We depicted this issue by comparing actual particle concentrations (measured by ICP-MS) to values obtained from MPS via calibration curves in water. We could detect a clear discrepancy for commercial PVP-coated Fe_3_O_4_, while especially SiO_2_-coated Zn-ferrites showed good agreement between measured and effective concentration.

This trend was verified by the obtained magnetic particle images and the used signals. SiO_2_-coated (70 wt%) Zn_0.4_Fe_2.6_O_4_ tracers could be clearly located in water and PBS, while PVP-coated Fe_3_O_4_ lost most of its signal in PBS and renders them inapplicable to MPI.

This highlights the importance of signal stability upon in vitro and in vivo exposure and the advantage of our prepared pre-aggregated Zn-ferrite tracers. The investigation of this issue during in vivo studies, as well as potential MPI signal evaluation approaches to reduce the effect, are topics for necessary research in the near future.

## Acknowledgements

We acknowledge Raphael Langenegger for the magnetic simulations. We thank the Swiss National Science Foundation for generous funding through the Eccellenza (181290) and the Scientific Exchange scheme (IZSEZ0_205894).

