## Supplementary information for "Particle interactions and their effect on magnetic particle imaging and spectroscopy"

### *Supporting Information*

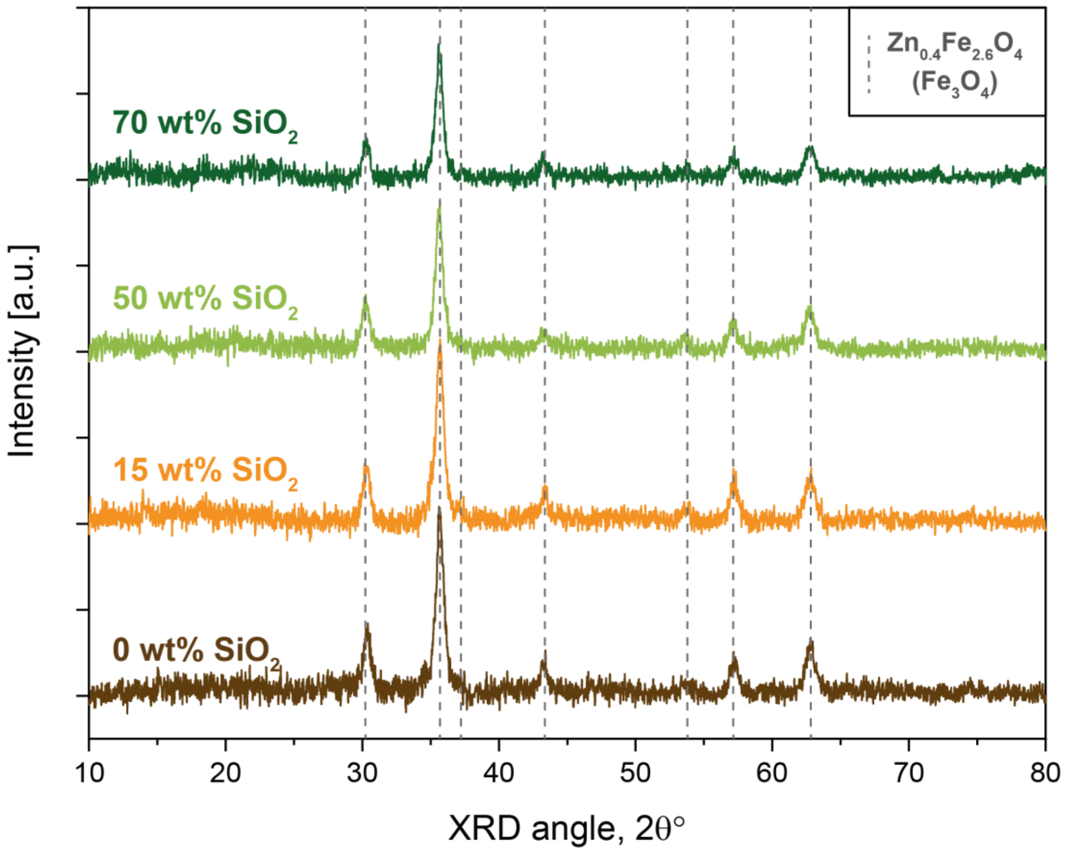


Figure S1. X-ray diffraction patterns of as-prepared $Zn_{0.4}Fe_{2.6}O_4$ nanoparticles with different amounts of $SiO_2$ coating. All particles show peaks characteristic to Zn-ferrites ($Fe_3O_4$).

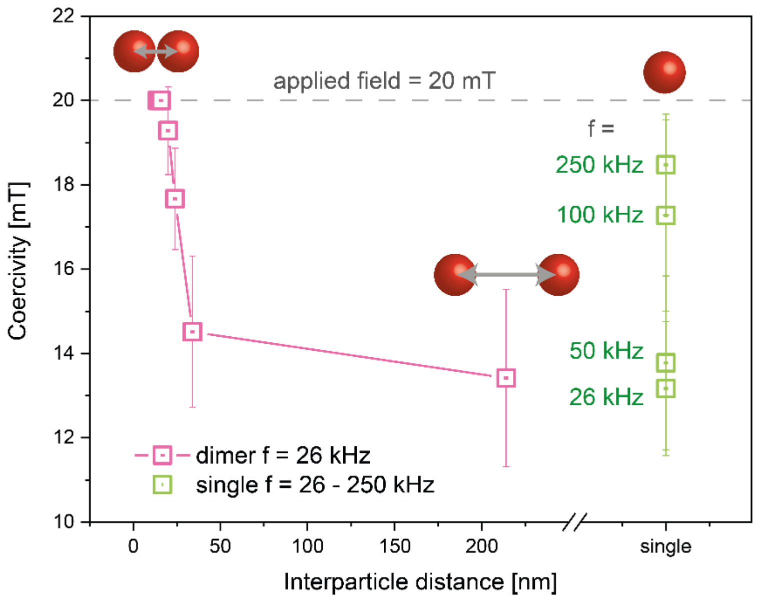


Figure S2. Values are determined from micromagnetic simulations. (Pink squares) Coercivities of dimers consisting of 14 nm $Fe_3O_4$ spheres as a function of their interparticle distance measured at applied field frequency of 26 kHz. (Green squares) Coercivities of a single particle as a function of applied field frequency. The dashed line indicates the maximum applied field strength (20 mT). For small interparticle distances, the applied field of 20 mT was not enough to reverse the magnetization. As a result, the coercivity remains at 20 mT up to a distance of 16 nm (2 nm gap). For larger distances, the coercivity decreases quickly and asymptotically reaches the value for a single particle of approximately 13 mT. Already at an interparticle distance of 34 nm (20 nm gap) $H_c$ is reduced to 14.5 mT. This trend shows the strong dependence of the magnetic properties on particle interactions, as previously discussed. These magnetic interactions, however, are distance-dependent and thus can be controlled by non-magnetic coatings, as shown above. The strong dependence on the frequency of the applied field is evident in the increased coercivity values for higher frequencies. As the field oscillates quicker, the magnetic dipoles of the nanoparticles have less time to align, which causes increased hysteresis. This is also detectable in the drastically lower coercivity values (< 1 mT) for the prepared bare and $SiO_2$-coated $Zn_{0.4}Fe_{2.6}O_4$ system measured in the quasi-static VSM setup (Table 1). Micromagnetic simulations were conducted via the Object Oriented MicroMagnetic Framework (OOMMF). A cellsize of 2 nm, crystalline anisotropy constant of 1.25 x $10^4$ J/m$^3$, exchange coefficient of 2.64 x $10^{11}$ J/m, temperature of 300 K, time steps of $10^{-12}$ s, saturation magnetization of 367709 A/m, gyromagnetic ratio of 2.21 x $10^5$ m/(A s) and damping coefficient α of 1 were used.

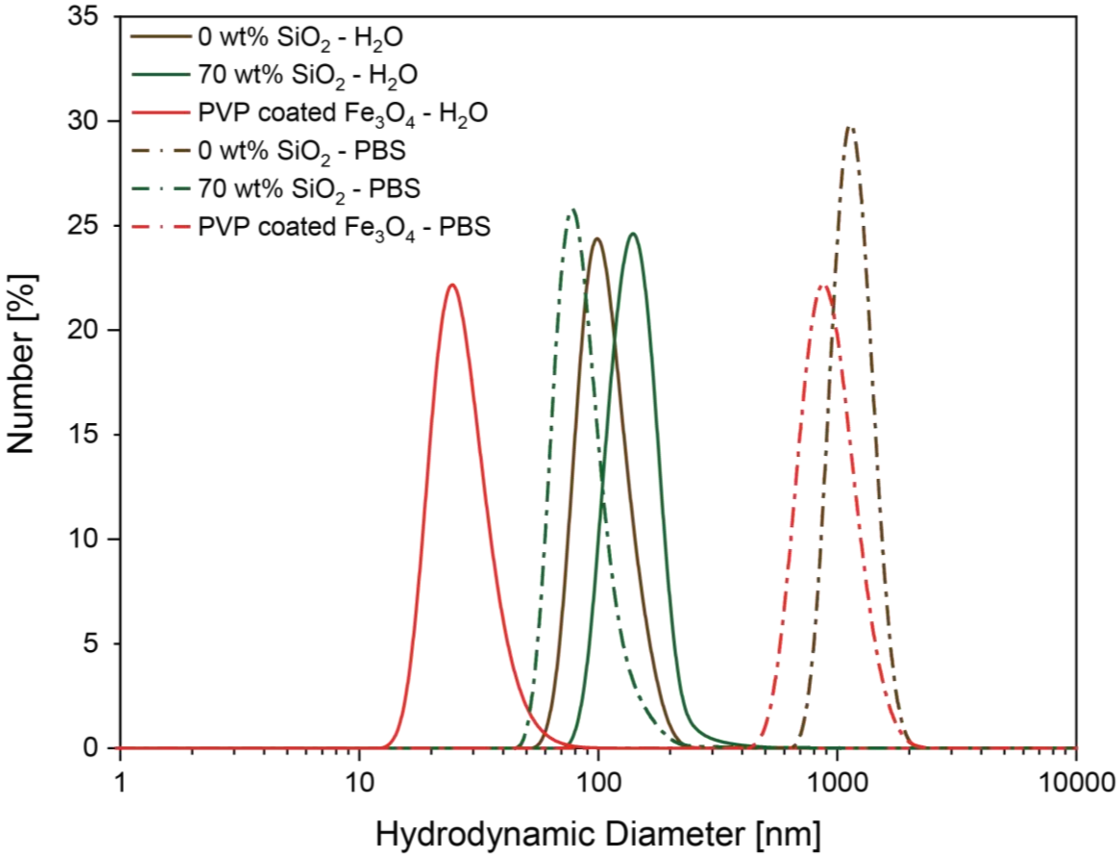


Figure S3. Number-weighted distributions of hydrodynamic diameters bare and $SiO_2$-coated (70 wt%) $Zn_{0.4}Fe_{2.6}O_4$ as well as of PVP-coated $Fe_3O_4$ in water and PBS. Measured immediately after sonication at a concentration of 0.1 mg/mL.
